# Cellular precision of orientation and spatial frequency maps in macaque V1

**DOI:** 10.1101/631416

**Authors:** Nian-Sheng Ju, Shu-Chen Guan, Shi-Ming Tang, Cong Yu

## Abstract

Functional organization of neuronal response properties along the surface of the neocortex is a fundamental guiding principle of neural computation in the brain. Despite this importance, the cellular precision of functional maps is still largely unknown. We address the challenge by using two-photon calcium imaging to measure cell-specific orientation and spatial frequency (SF) responses across fields of macaque V1 superficial layers. The cellular orientation maps confirm iso-orientation domains, but rarely show pinwheels. Pinwheels obtained through conventional Gaussian smoothing and vector summation of orientation responses mostly overlap with blood vessel regions, suggesting false singularities. Cellular SF maps clarify existing controversies by showing weak iso-frequency clusters, which also suggests a weak geometric relationship between orientation and SF maps. Most neurons are tuned to medium frequencies, but the tuning functions are often asymmetric with a wider low- or high-frequency branch, which may help encode low or high SF information for later decoding.

## Introduction

Neurons in primate V1 are functionally organized according to their preferences to stimulus features (Hubel & Wiesel, 1962, 1968), likely a result of wiring optimization (Chklovskii & Koulakov, 2004) or self-organization during evolvement (Kaschube et al., 2010). The local structures of functional maps may be correlated to the tuning properties of neurons (Callaway, 1998; McLaughlin et al., 2003; Nauhaus et al., 2008), such that V1 neurons near the orientation pinwheels tend to have broader tuning than those in iso-orientation domains (Nauhaus et al., 2008). These correlations may suggest the guiding roles of functional organization in neural computation.

Orientation and spatial frequency are two fundamental stimulus features. It is well known that the orientation preferences of V1 neurons are mapped in iso-orientation domains that vary smoothly (Hubel & Wiesel, 1962, 1963), and that different iso-orientation domains converge at singularity points or pinwheels (Bonhoeffer & Grinvald, 1991). Nevertheless, Hubel and Wiesel (2005) disagreed on the idea of pinwheels from optical imaging experiments because pinwheels are not centered on blobs where neurons have poor orientation tuning, likely an artifact caused by the limited resolution of optical imaging (Polimeni et al., 2005).

V1 spatial frequency maps are less understood and somewhat controversial (Issa et al., 2008; Nauhaus et al., 2012). Single-unit results from monkey V1 and cat area 17 are conflicting on whether spatial frequency preferences change smoothly or abruptly (Silverman et al., 1989; Born & Tootell, 1991; DeAngelis et al., 1999; Molotchnikoff et al., 2007). More recent optical imaging results tend to show clusters of spatial frequency tuning, but are inconsistent on whether these clusters are continuous, pinwheeled, or not repeating at all (Everson et al., 1998; Issa et al., 2000; Xu et al., 2007). To complicate the matter, there is suspicion that the spatial frequency maps from intrinsic optical imaging signals may have been greatly influenced by vascular artifacts, which may be mistakenly interpreted as spatial frequency domains (Sirovich & Uglesich, 2004). Such vascular artifacts should apply to orientation maps too.

Single-unit recording data may be inaccurate due to under-sampling. Intrinsic optical imaging, on the other hand, lacks cellular resolution. The two-photon imaging technology is able to overcome these difficulties in large measure by simultaneously recording hundreds of neurons’ activities at single cell precision (Ohki et al., 2005; Nauhaus et al., 2012; Nauhaus et al., 2016; Li et al., 2017). Most relevant to the current study, Nauhaus et al. (2012; 2016) have studied V1 orientation and spatial frequency tuning in anesthetized macaques, using two-photon imaging with a bulk-loaded calcium indicator Oregon Green BAPTA-1 AM (OGB-1). They obtained orientation maps with iso-orientation domains radiating from pinwheels, as well as spatial frequency maps with continuous iso-spatial frequency domains. However, these maps on the basis of “large-scale” imaging are still pixel based and lack single cell details. That is, the orientation and spatial frequency maps show each pixel’s, not each cell’s, orientation and spatial frequency preferences, respectively. Here the orientation preference of a specific pixel is the vector sum of the pixel’s responses to 8 orientations, and the spatial frequency preference is the center-of-mass of the pixel’s tuning function measured with 6 spatial frequencies.

We decided to use long-term two-photon calcium imaging (Li et al., 2017) to study cellular orientation and spatial frequency maps in awake macaque monkeys. Our setup allowed us to simultaneously record hundreds of neurons’ responses at single neuron resolution in an imaging window of 850 × 850 μm^2^, which was large enough to reveal the two-dimensional layouts of orientation and spatial frequency preferences, and was similar in size to the large-scale imaging window of Nauhaus et al. (2012; 2016). Unlike the short imagining time when OGB-1 is used as the calcium indicator, the calcium indicator GCaMP5 makes it possible to image the same neurons’ activities for an extended period (Li et al., 2017). For each imaging window we were able to measure at two recording depths (150 and 300μm) the neuronal responses to a Gabor stimulus at 12 orientations, 6 spatial frequencies, 2 opposite drafting directions, and 3 sizes (to maximize summation and minimize surround suppression), to achieve detailed response mapping. We also used 1∼1.5 s inter-stimulus intervals to minimize the interferences of responses from previous stimulus presentations.

## Results

We recorded orientation and spatial frequency responses in V1 superficial layers II/III in four awake macaque monkeys (Figure 1a). Recordings were made at 6 locations (monkeys A, C each with two nearby locations, and monkeys B, D each with one location) at depths of 150 and 300 μm from the cortical surface, so that 12 sets of data were collected. Initial screening with bandpass filtering and thresholding of differential images (ΔF = F - F0) (see Methods) identified 12534 regions of interests (ROIs) or possible cell bodies (Figure 1b). Each cell’s responses (ΔF/F0) were then fitted with a Gaussian model to estimate orientation tuning, and a Difference-of-Gaussian model to estimate spatial frequency tuning. A total of 11225 cells (89.6%) were tuned to orientation and/or spatial frequency, and were included in data analysis (examples shown in Figure 1d, e). Among them, 6874 (61.2%) were tuned to both orientation and spatial frequency, 1924 (17.1%) to orientation only, and 2427 (21.6%) to spatial frequency only.

**Figure 1.**
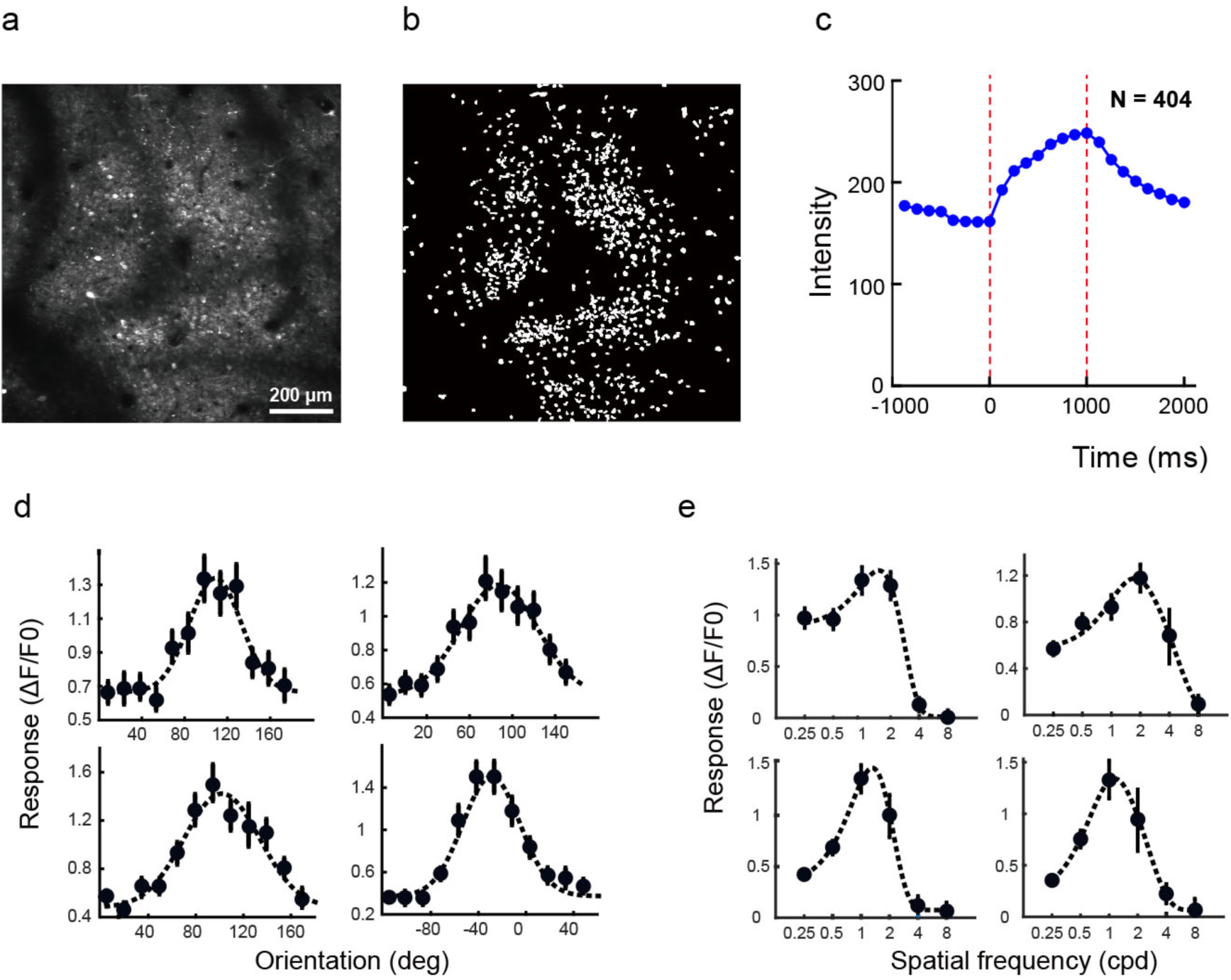
Two-photon calcium imaging and orientation and spatial frequency tuning functions. **a**. The average two-photon image after alignment correction over a recording session at Monkey A recording site 1 at 150 μm depth. **b**. Regions of interests (ROIs) or cell bodies extracted from **a** after band-pass filtering and thresholding (>= 3 standard deviation of the mean). **c**. The mean time course of calcium responses over all neurons that were tuned to both orientation and spatial frequency at monkey A site 1 at 150 μm depth, each at the optimal stimulus orientation and spatial frequency. Two vertical dashed lines indicate stimulus onset and offset. Each dot indicates one frame (8 frames per second, 125 ms per frame). The mean response started to saturate around 5th-6th frames (625-750 ms) after stimulus onset. **d**. Examples of individual orientation tuning functions fitted with a Gaussian model. **e**. Examples of individual spatial frequency tuning functions fitted with a Difference-of-Gaussian model. Error bars represent ±1 SEM.

### Cellular orientation maps

Cellular orientation maps contained various sizes and shapes of patches separated by blood vessels (Figure 2a). Each patch contained clear clusters of neurons tuned to similar orientations. These iso-orientation domains changed preferred orientations smoothly. Moreover, the distributions of orientations domains appeared similar at 150 and 300 μm depths, suggesting vertical orientation columns. These results confirmed previous reports of iso-orientation domains and orientation columns (Hubel & Wiesel, 1962; Bonhoeffer & Grinvald, 1991).

**Figure 2.**
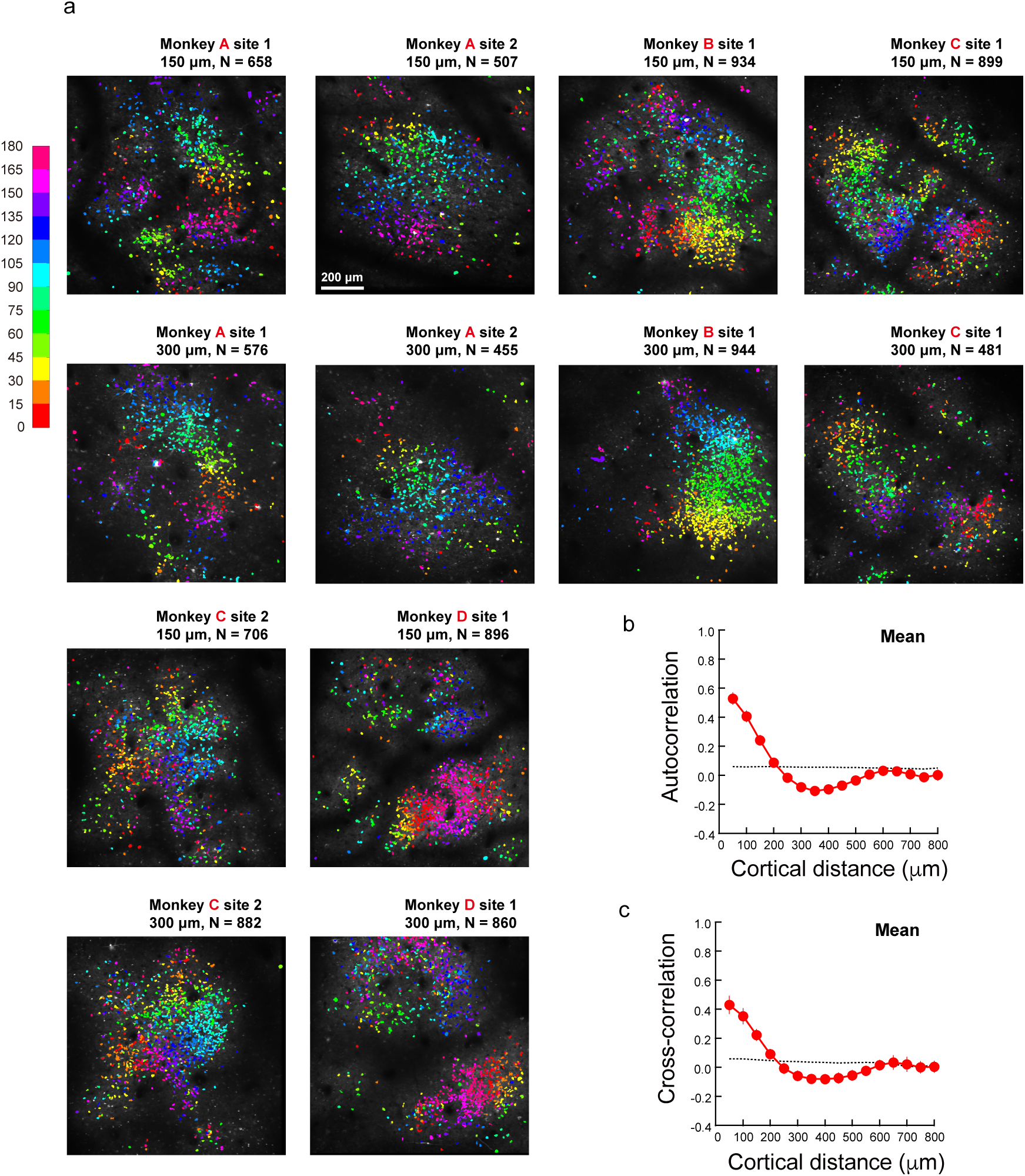
Cellular orientation maps. **a.** Cellular orientation maps from 4 macaques at two recording depths. **b**. Mean within-map autocorrelation of orientation tuning over 12 orientation maps as a function of the absolute cortical distance. Each red dot indicates the averaged measured autocorrelation within a 50 μm bin up to the dot’s corresponding cortical distance on the x-axis. The black dashed curve is the baseline autocorrelation simulated with neurons randomly shuffled in positions. **c.** Mean cross-correlation of orientation tuning between two depths over 6 recording sites as a function of the absolute cortical distance. Each red dot indicates the averaged measured cross-correlation within a 50 μm bin up to the dot’s corresponding cortical distance on the x-axis. The black dashed curve represents the mean baseline cross-correlation. Error bars represent ±1 SEM.

To quantify the iso-orientation domains, we calculated the autocorrelations of orientation tuning as a function of absolute cortical distance within the same map. Compared to the mean baseline autocorrelations with neurons randomly shuffled (mean = 0.053 within 800 μm), the measured mean autocorrelations over 12 orientation maps were significantly higher within cortical distances of 0-50 μm, 50-100μm, and 100-150 μm (p_s_ < 0.001, two tailed paired t-tests here and later), and became similar within 150-200 μm (p = 0.133) (Figure 2b). The normalized (measured/baseline) autocorrelations were 8.86, 7.14, and 4.04 within 0-50, 50-100, and 100-150 μm, respectively. Data from individual maps were quite similar to the mean. These results suggest significant iso-orientation domains that were about 150 μm in width.

And to quantify the columnar structures of orientation preferences, we calculated the cross-correlations of orientation tuning as a function of absolute cortical distance between maps at two depths, which showed a similar trend. The mean cross-correlations over all recording sites (0.43, 0.35, and 0.22 within 0-50, 50-100, and 100-150 μm, respectively) were 7.44, 6.19, 4.22 times as high as the corresponding baselines (mean = 0.056 across 3 distances) (p_s_ < 0.011), respectively, and became insignificantly different at 150-200 μm (p = 0.142) (Figure 2c). Individual maps showed similar trends, except for the one with monkey A site 2 that showed much poorer alignment between two depths (normalized autocorrelations = 1.67, 1.70, and 1.67 within 0-50, 50-100, and 100-150 μm, respectively). Therefore, V1 neurons in macaques were organized vertically in orientation columns, but with exceptions.

What really surprised us in these cellular orientation maps was the rare appearance of pinwheels despite the apparent orientation clusters. The distributions of cells in these maps were divided into patches by blood vessels (and likely shallows of more superficial blood vessels). Within each patch, iso-orientation domains rarely appeared to converge in pinwheels (Figure 2a). As pinwheel structures are mostly revealed by optical imaging, we constructed optical imaging-like orientation maps, in which the responses to each orientation were first Gaussian smoothed, and then the orientation preference of each pixel was independently calculated (Figure 3b, example maps of monkey A recording site 1; see Methods). The optical imaging-like orientation maps did show numerous pinwheels (e.g., 4 on the upper map and 5 on the lower map in Figure 3b). However, when cellular orientation maps (Figure 2a) and contours of optical imaging-like maps were overlaid (Figure 3c), it was apparent that all 9 pinwheels were located in regions with few neurons recorded. As more easily discernable in Figure 3a, these pinwheels were either overlaid with radial or vertical blood vessels, or in the shallows of likely more superficial blood vessels.

**Figure 3.**
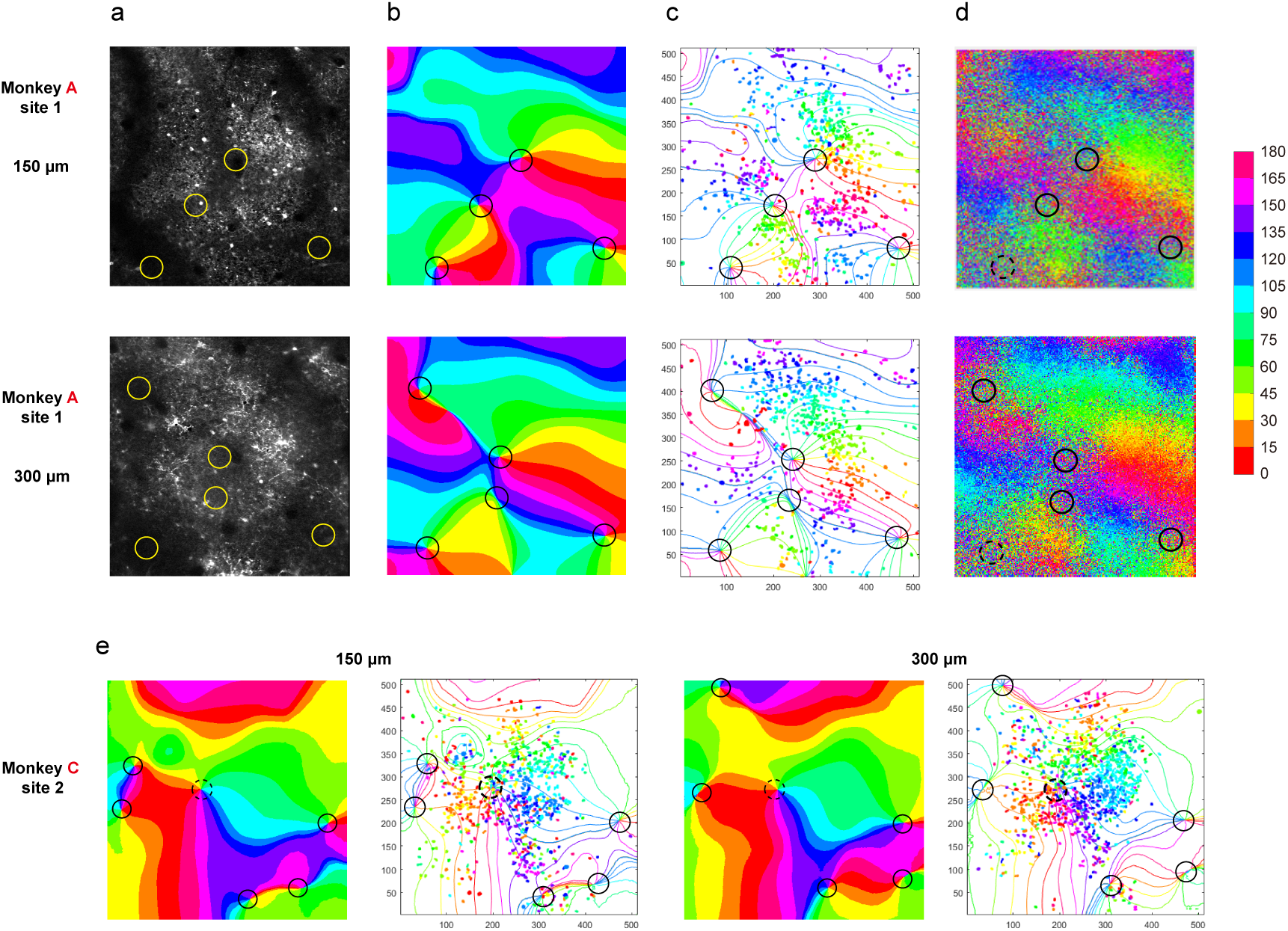
Comparisons of cellular and optical imaging-like orientation maps. **a.** The two-photon images over a recording session at Monkey A recording site 1 at 150 and 300 μm depths. The circles in each map indicate locations of pinwheels revealed from optical imaging-like orientation maps in **b**. **b**. Optical imaging-like orientation maps with the data sets of monkey A site 1. **c**. Overlaid cellular orientation maps of monkey A site 1 from Fig. 2a and contoured optical imaging-like orientation maps from **b**. **d**. Orientation maps obtained using the methods of Nauhaus et al. (2012; 2016). The same circles in each corresponding map indicate locations of pinwheels in optical imaging-like maps from **b**. **e**. Optical imaging-like orientation maps and overlaid cellular and contoured optical imaging-like orientation maps of monkey C site 2 at 150 μm (left two panels) and 300 μm (right two panels). The maps at each recording depth contain a possible real pinwheel as indicated by a dashed circle.

Pinwheels were also evident in orientation maps through large-scale two-photon imaging (Nauhaus et al., 2012; 2016). These maps were different from our optical imaging-like maps (Figure 3b) in that each pixel’s orientation preference was calculated without initial Gaussian smoothing of responses to each orientation. We followed Nauhaus et al. (2012; 2016) to recreate orientation maps using data sets from monkey A site 1 (Figure 3d). These orientation maps did not show clear boundaries of iso-orientation domains as did maps in Nauhaus et al. (2012; 2016), but at least some of regions with possible pinwheels (e.g., solid circles) appeared to correspond to the pinwheels estimated in Figure 3b, and most of them were actually located in blood-vessel regions.

Overall, among 50 pinwheels revealed in all 12 optical imaging-like orientation maps, only 2 were likely real pinwheels (indicated by dashing circles in maps of monkey C site 2, Figure 3e). All others were located in regions with blood vessels or with few neurons. These results suggest possible false singularities or pinwheel centers in optical imaging-like orientation maps.

### Cellular spatial frequency Maps

The cellular spatial frequency maps showed poor iso-frequency domains (Figure 4a). The mean within-map autocorrelation over 12 maps was 0.67, 0.63, and 0.59 within 0-50, 50-100, and 100-150 μm, respectively, slightly but significantly higher than the corresponding baselines, all of which at 0.52 (p < 0.001), and became insignificantly different within 150-200 μm (p = 0.114) (Figure 4b). The normalized autocorrelations were 1.28, 1.20, and 1.12 within 0-50, 50-100, and 100-150 μm, respectively, which were close to the chance level at 1, in sharp contrast to high orientation autocorrelations at 8.86, 7.14, and 4.04 within corresponding cortical distances.

**Figure 4.**
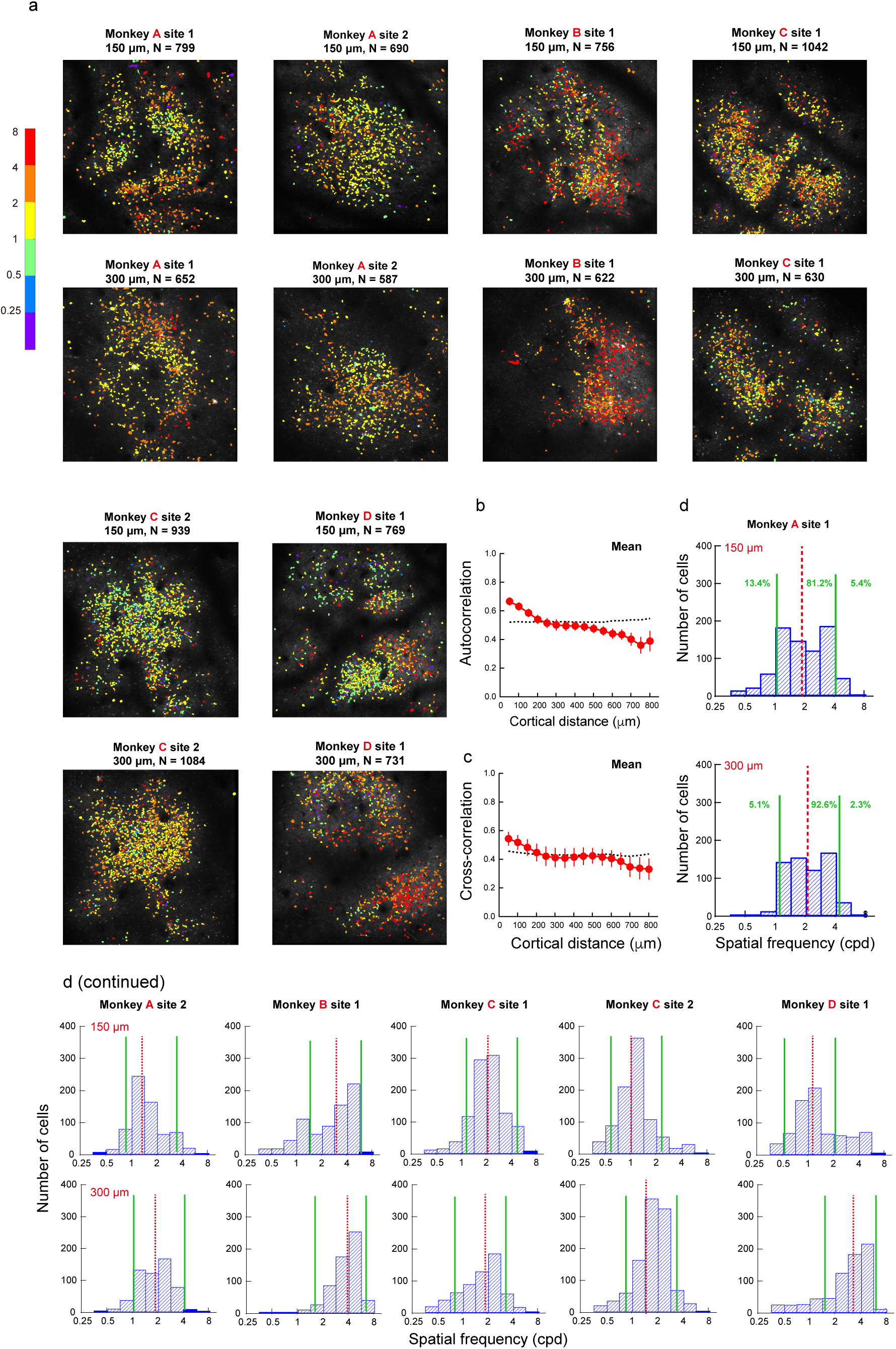
Cellular spatial frequency maps. **a**. Cellular spatial frequency maps from 4 macaques at two recording depths. **b**. Mean within-map autocorrelation over 12 spatial frequency maps as a function of the absolute cortical distance. Each red dot indicates the averaged measured autocorrelation within a 50 μm bin up to the dot’s corresponding cortical distance on the x-axis. The black dashed curve represents the baseline autocorrelation with shuffled neurons. **c.** Mean cross-correlation of spatial frequency maps between two depths over 6 recording sites as a function of the absolute cortical distance. Each red dot indicates the averaged measured cross-correlation within a 50 μm bin up to the dot’s corresponding cortical distance on the x-axis. The black dashed curve represents the baseline cross-correlation with shuffled neurons. **d**. The distributions of neurons against preferred spatial frequency in half-octave steps in monkey A site 1 at 150 and 300 μm depths. The pair of green vertical solid lines indicate the low and high boundaries of a 2-octave range of preferred spatial frequencies. The red vertical dashed line indicates the median preferred spatial frequencies.

Cross-correlations of spatial frequency tuning between two depths were also very poor even within the first 100 μm of the cortical distance (Figure 4c). The mean measured vs. baseline cross-correlations was 0.55 vs. 0.46 (p = 0.033) and 0.52 vs. 0.45 (p = 0.033) within 0-50 and 50-100 μm, respectively, with normalized cross-correlations at 1.20 and 1.16, much lower than the orientation cross-correlations of 7.44 and 6.19 within corresponding cortical distances. The difference became insignificant at 100-150 μm (p = 0.070). These data indicated weak columnar structures in spatial frequency maps.

The low normalized autocorrelations and cross-correlations could be mainly attributed to baseline autocorrelations and cross-correlations that were about 10 times or more as high than those with orientation maps (0.53 vs. 0.05 with mean autocorrelation and 0.44 vs. 0.03 with mean cross-correlation across all distances up to 800 μm). The high baselines mostly resulted from a narrow range of preferred spatial frequencies in each map. Take the maps of monkey A site 1 for example (Figure 4d), 81.2% neurons’ preferred spatial frequencies fell into a 2-octave range from 1.01 to 4.04 cpd at 150 μm, and 92.6% fell into a 2-octave range from 1.07 to 4.41 cpd at 300 μm. Moreover, there were far less neurons tuned to spatial frequencies lower than 1 cpd or higher than 4 cpd. Other spatial frequency maps also showed similar narrow ranges of preferred spatial frequencies and scarcity of cells tuned to low or high spatial frequencies (Figure 4d). On the average 83.0% of neurons from all monkeys were tuned to a 2-octave range of spatial frequencies, with 10.6% tuned to lower frequencies and 6.4% to higher frequencies. These trends were largely consistent with De Valois et al. (1982) who showed that most V1 neurons’ peak frequencies spread over a 2-octave range (approximately 1.4-5.6 cpd for Y cells and 1.0-4.0 cpd for X cells) at the comparable eccentricities (3-5°) in anesthetized macaques. The exact ranges of the 2-octave range varied among 12 maps. Many were around 1-4 cpd, but a few were either 0.5-1 octaves lower (monkey A site 2, monkey C site 2, and monkey D site 1, all at 150 μm; Figure 4d) or higher (monkey B at two depths and monkey D at 300 μm; Figure 4d). The shifts of spatial frequency ranges were not related to retinal eccentricity. For example, the recording sites of monkeys B and D were not closer to the fovea than the other two monkeys. Rather the shifts might suggest individual differences, or clusters of spatial frequency tuning possibly in a grander scale than orientation clusters. More data are required to verify these possibilities. Nevertheless, none of these maps contained many neurons tuned to spatial frequencies lower than 0.5 cpd or higher than 8 cpd.

Behaviorally, macaque monkeys, like humans, are sensitive to low spatial frequencies, as well as to high spatial frequencies up to ∼30 cpd at comparable retinal eccentricities (De Valois et al., 1974). How could low and high spatial frequency information be represented in the brain if there are too few V1 neurons tuned to them? On possible strategy is to have these low and high spatial frequencies first encoded by medium-frequency V1 neurons, and then decoded from population responses by downstream neurons. For example, assuming the simplest case in which two neurons tuned to the same medium frequency but have different slopes of low (or high) frequency branches, subtracting two tuning functions would give a differential function that contains a lower (or higher) spatial frequency peak.

We calculated the lower- and higher-half bandwidths at half height for the spatial frequency tuning function of each neuron in our dataset, and contrasted the higher-half bandwidth against the lower-half bandwidth for all neurons at 150 and 300 μm depths (Figure 5). The median lower-half bandwidths were 1.17 octaves at 150 μm and 1.16 octaves at 300 μm, about 0.4 octaves wider than the median higher-half bandwidths that were 0.77 octaves at both depths. Therefore, more neurons’ spatial frequency tuning functions contained a wider low frequency branch that would encode low frequency information, like those shown in Figure 1e. Still, there were also many neurons whose tuning functions contained a wider high frequency branch to encode high spatial frequency information. Some neurons had a very wider lower-half or higher-half bandwidth of >2 octaves. These were mostly low-pass or high-pass neurons that would encode very low or very high (up to cutoff) spatial frequencies.

**Figure 5.**
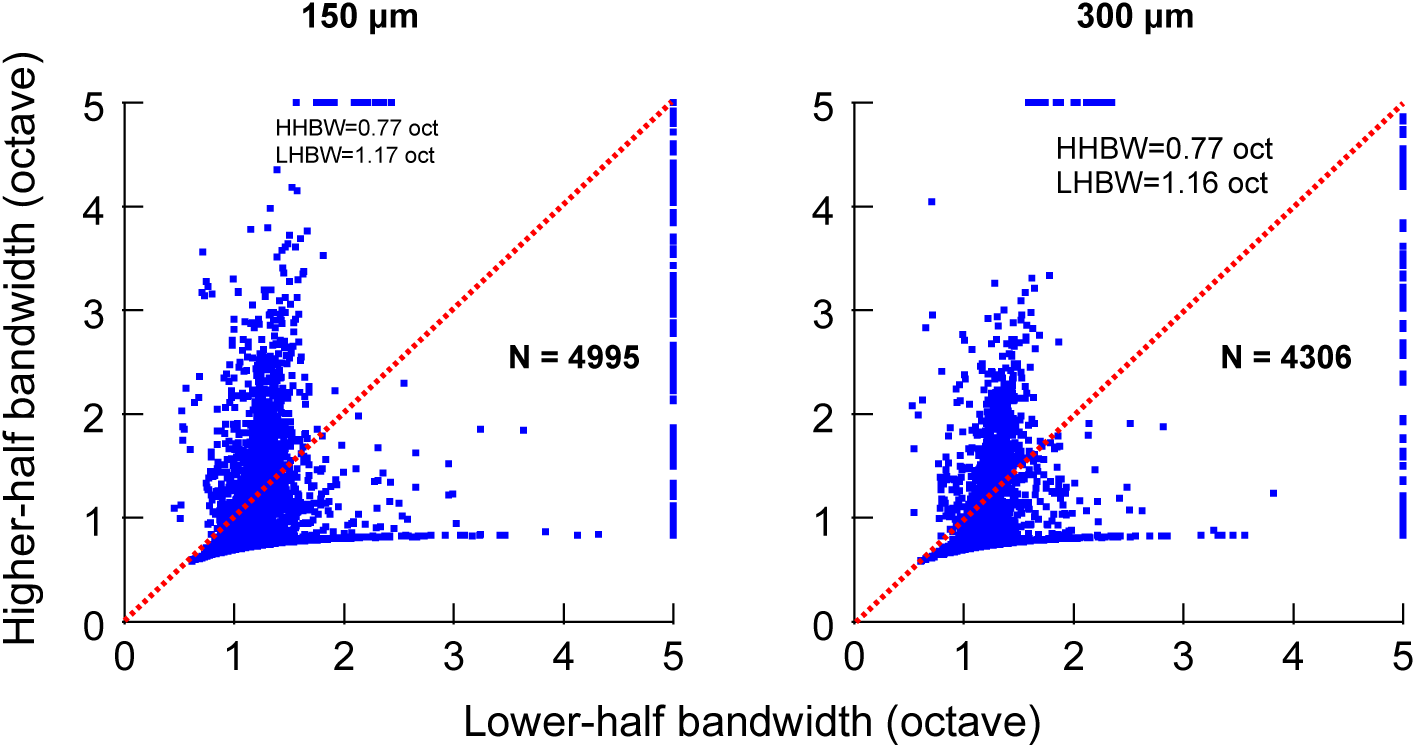
All neuron’s lower-half bandwidths (LHBW) and higher-half bandwidths (HHBW) of spatial frequency tuning functions are contrasted at two recording depths. A low-pass neuron’s LHBW is assigned to be 5 if not computable. So is a high-pass neuron’s HHBW. Error bars represent ±1 SEM.

## Discussion

In this study we used two-photon imaging with genetically encoded calcium indicators (GCaMP5) to study cellular orientation and spatial frequency maps in awake macaque V1. Cellular orientation maps confirm iso-orientation domains and orientation columns, but reveal scarce evidence for pinwheels. The orientation maps are divided by blood vessels into various sizes and shapes of patches, in which the iso-orientation domains rarely converge on singularity points or pinwheels. Instead, many pinwheels revealed in optical imaging-like orientation maps through Gaussian-smoothing and vector summation of orientation responses tend to be centered on neuron-less blood vessel regions. Although we are not questioning the very existence of orientation pinwheels, indeed there are two likely pinwheels in monkey C (Fig. 3e), as well as in previous two-photon imaging studies (Ohki et al., 2005; Li et al., 2017), our results suggest that real pinwheels could be accidental occurrences in V1 orientation maps, at least at superficial layers where our recordings were performed.

The cellular spatial frequency maps reveal weak iso-spatial frequency domains and columnar structures. This finding is consistent with Sirovich and Uglesich (2004) that V1 spatial frequency maps from optical imaging contain no columnar structures after removing the vascular effects. After Gaussian-smoothing and vector summation of the raw spatial frequency responses, we did find in optical imaging-like spatial frequency maps that the blood vessel regions were filled with colors of nearby neurons to form false iso-frequency domains. The findings that most neurons are tuned to a narrow range of medium spatial frequencies (∼2 octaves), and only a small percentage of neurons are tuned to low and high spatial frequencies, are largely consistent with De Valois et al. (1982), as we mentioned earlier, as well as Edwards et al. (1995) who reported that blob neurons and most interblob neurons in macaque V1 share similar preferred medium spatial frequencies with a low boundary of 1.4 cpd, except that some interblob neurons are tuned to higher spatial frequencies. In addition, the narrow range of spatial frequency tuning agrees with a psychophysical masking study that spatial frequency tuning of spatial channels is limited within a 2-octave range (Kontsevich & Tyler, 2013).

Nevertheless, human and monkeys are sensitive to low and high spatial frequencies. This dilemma could be solved by many middle frequency neurons whose tuning functions either a wider low-frequency branch or a wider high-frequency branch to encode low or high frequency information. Foster et al. (1985) reported in a single-unit study that V2 neurons tend to be tuned to spatial frequencies 2-octave lower than V1 neurons do. At comparable retinal eccentricities (2-5°), the preferred spatial frequencies range from 0.5 to 8.0 cpd in V1, similar to our data, and are about two octaves lower from 0.2 to 2.1 cpd in V2. It is likely that low spatial frequency information is first encoded by medium frequency V1 neurons, and then decoded by low-frequency V2 neurons. However, it is unclear where high spatial frequency information could be decoded.

It is imperative to compare our two-photon imaging results and those of Nauhaus et al. (2012; 2016), the only two-photon imaging studies that constructed orientation and spatial frequency maps in macaque V1. Nauhaus et al. (2012; 2016) relied on “large-scale” pixel-based imaging to obtain orientation and spatial frequency maps, and “fine-scale” imaging to measure tuning functions of individual neurons and calculate pairwise correlations. Fine-scale imaging only imaged a 200 × 200 μm^2^ area with a relatively small number of neurons. We were able to obtain cellular orientation and spatial frequency maps from similar-sized imaging windows (850 × 850 μm^2^), but with doubled resolution (1.6 vs. 3.0 μm pixel^-1^) and a lot more neurons. For example, the total number of neurons considered in analysis by Nauhaus et al. (2012) was 735, in contrast to over 11,000 orientation and/or spatial frequency tuned neurons in our study.

### Orientation maps

The pixel-based orientation maps in Nauhaus et al. (2012; 2016) show iso-orientation domains and pinwheels. Our cellular orientations maps, each divided by blood vessels into various sizes and shapes of patches, also reveal iso-orientation domains, but rarely pinwheels. Instead optical imaging-like orientation maps through Gaussian smoothing and vector summation of raw orientation responses led to many false pinwheels located on neuron-less blood vessel regions. The blood-vessel artifacts were not considered by Nauhaus et al. (2012; 2016).

Autocorrelation indicates the degree of functional clustering. Nauhaus et al. (2012) reported that normalized orientation autocorrelation declines gradually from ∼8 to ∼1 within a cortical distance from 7-52 μm to around 200 μm. Similarly, our data show a decline from ∼9 to ∼1 within comparable ranges of cortical distances. Nauhaus et al.’s autocorrelations were estimated from the entire data set from 10 images, with the assumption that neurons in these images together represent an unbiased sample. Our estimates were first obtained from each of 12 images and then averaged.

### Spatial frequency maps

Hauhaus et al. (2012) found that within cortical distances of 7-52 mm, the normalized autocorrelation is ∼3, weaker than ∼8 with orientation maps. We also found weaker spatial frequency clustering with a similar periodicity, but the clustering is even weaker, as the normalized autocorrelation is merely 1.28 within a cortical distance of 50 mm. Hauhaus et al.’s (2012) maps have a wider range of spatial frequency preferences than ours. The spatial frequency maps from two studies both have a similar low boundary of preferred spatial frequencies at 0.5-1 cpd, but Nauhaus et al.’s maps include preferences to higher spatial frequencies at 4-8 cpd. Some of our maps do show tuning to higher spatial frequencies, but the ranges of spatial frequency tuning are not getting wider, but also shifting to higher frequencies (Figure 4a). As a main conclusion, Nauhaus et al. (2012) reported that orientation and spatial frequency maps are orthogonal. However, such a relationship is difficult to demonstrate in our cellular maps because of the very weak iso-spatial frequency domains.

In summary, our cellular orientation maps reveal iso-orientation domains and columnar structures, but hardly pinwheels, in superficial layers of macaque V1. Many pinwheels revealed by optical imaging-like orientation maps actually result from false singularities because of the blood vessel effects, suggesting that functional micro-organizations revealed with non-cellular level techniques need to be treated with great caution. Cellular spatial frequency maps show poor functional organization of spatial frequency tuning, largely because most neurons prefer a limited range of medium frequencies and are spatially intermixed. However, most of these neurons have asymmetric tuning functions that contain a wider branch toward either low or high spatial frequencies, which may be primarily responsible for encoding low or high spatial frequency information for later decoding.

## Methods

### Monkey preparation

Four macaque monkeys (aged 5-8 years) were used in this study. Each monkey was prepared with two sequential surgeries under general anesthesia and strictly sterile conditions. In the first surgery, a 20-mm diameter craniotomy was performed on the skull over V1. The dura was opened and 100-150 nL AAV1.hSynap.GCaMP5G.WPRE.SV40 (AV-1-PV2478, titer 2.37e13 (GC/ml), Penn Vector Core) was pressure-injected at a depth of ∼350 μm. The dura was then sutured, the skull cap was re-attached with three titanium lugs and six screws, and the scalp was sewn up. After the surgery, the animal was returned to the cage, treated with injectable antibiotics (Ceftriaxone sodium, Youcare Pharmaceutical Group, China) for one week. The second surgery was performed 45 days later. A T-shaped steel frame was installed for head stabilization, and an optical window was inserted onto the cortical surface. More details of the preparation and surgical procedures can be found in Li et al. (2017). The procedures were approved by the Institutional Animal Care and Use Committee at Beijing University.

### Behavioral task

After a ten-day recovery from the second surgery, monkeys were seated in primate chairs with head restraint. They were trained to hold fixation on a small white spot (0.1°) with eye positions monitored by an ISCAN ETL-200 infrared eye-tracking system (ISCAN Inc.) at a 500-Hz sampling rate. During the experiment, a trial with the eye position deviated 2° from the fixation point was discarded and repeated. For the remaining trials, the eye positions were mostly concentrated around the fixation point. The mean and standard deviation of the eye positions, as well as the ratio of trials with eye positions within 0.5° from the fixation point, were 0.17°, 0.12°, and 98.6% for monkey A; 0.23°, 0.29°, and 97.1% for monkey B; 0.22°, 0.12°, and 98.4% for monkey C; and 0.16°, 0.28°, and 97.3% for monkey D.

### Visual stimuli

Visual stimuli were generated by the ViSaGe system (Cambridge Research Systems) and presented on a 21’’ CRT monitor with a refresh rate of 80 Hz. Monitor resolution was set at 1280 pixel × 960 pixel. Because of the space and monitor size limits, viewing distances varied depending on the stimulus spatial frequency (30 cm at 0.25/0.5/1cpd, 60 cm at 2cpd, and 120 cm at 4/8 cpd). Imaging of all orientations and spatial frequencies at a specific recording depth was completed within a session that lasted for 3-4 hours.

A drifting square-wave grating (full contrast, 4 cpd spatial frequency, 3 cps temporal frequency, and 0.4° diameter) was first used to determine the receptive field location of each imaging window, which was around 2-4° eccentricity. For orientation and spatial frequency tuning measurements, the stimulus was a high-contrast (0.9) Gabor gating (Gaussian-windowed sinusoidal grating) drifting at 2 cycles/s in opposite directions perpendicular to the Gabor orientation. The Gabor grating varied at 12 equal-spaced orientations from 0° to 165° in 15° steps, and 6 spatial frequencies from 0.25 to 8 cpd in 1 octave steps. It also varied in 3 sizes at approximately 1, 1.5, and 2 octaves (or σ = 0.42λ, 0.64λ, and 0.85λ) at 0.25, 0.5, and 1 cpd, and 1.5, 2, and 2.5 octaves (or σ = 0.64λ, 0.85λ, and 1.06λ) at 2, 4, and 8 cpd, respectively (λ = wavelength). Our pilot measurements found very strong surround suppression with larger stimuli. Therefore, we decided to use these three stimulus sizes (always >= 1 octave) at each spatial frequency, so that the best responses of each neuron could be roughly estimated with the most center excitation and least surround suppression.

The stimuli at a specific viewing distance were pseudo-randomly presented. Each stimulus was presented for 1 s, with an inter-stimulus interval of 1 s (monkeys A, B) or s (monkey C, D) to minimize the interference of responses from previous trials (Fig. 1c). Each stimulus condition was repeated 10 times for monkeys A, B and 12 times for monkeys C, D, with half trials for each opposite direction.

### Two-photon imaging

Two-photon imaging was performed with a Prairie Ultima IV (In Vivo) two-photon microscope (Prairie Technologies) and a Ti:sapphire laser (Mai Tai eHP, Spectra Physics). One or two windows of 850 × 850 μm^2^ were selected in each animal and imaged using 1000-nm femtosecond laser under a 16 × objective lens (0.8 N.A., Nikon) at a resolution of 1.6 μm/pixel. Fast resonant scanning mode (32 fps) was chosen to obtain continuous images of neuronal activity (8 fps after averaging every 4 frames).

### Imaging data analysis

Data were analyzed with customized MATLAB codes. A normalized cross-correlation based translation algorithm was used to reduce motion artifacts (Li et al., 2017). Specifically, a template image was first generated by averaging 1000 successive frames in one imaging session. Two-photon images of one cortical area across days were then corrected and aligned using the template image. After the correction, fluorescence changes were associated with corresponding visual stimuli through the time sequence information recorded by Neural Signal Processor (Cerebus system, Blackrock Microsystem). By subtracting the mean of the 4 frames before stimuli onset (*F0*) from the average of the 6th-9th frames after stimuli onset (*F*) across 5 or 6 repeated trials for the same stimulus condition (same orientation, spatial frequency, size, and drifting direction), the differential images (*ΔF = F - F0*) were obtained. These differential images were then filtered with a band-pass Gaussian filter (size = 10-20 pixels). Finally, connected subsets of pixels (>25 pixels) with average pixel value > 3 standard deviations of the mean brightness were classified as regions of interests (ROIs) or potential neurons.

Once the ROIs or potential neurons were decided, the ratio of fluorescence change (*ΔF/F0*) was calculated as neuronal responses. Each stimulus condition was presented for 10-12 trials, half for each of two opposite directions. For a specific cell’s response to a specific stimulus condition, the *F0*_*n*_ of the n-th trial was the average of 4 frames before stimulus onset, and F_n_ was the average of 5th-8th or 6th-9th frames after stimulus onset, whichever was greater. F0_n_ was then averaged across 10 or 12 trials to obtain the baseline F0 for all trials (for the purpose of reducing variations), and ΔF_n_/F0 = (F_n_-F0)/F0 was taken as the neuron’s response to this stimulus at this trial. A small portion (6.3%) of the neurons showed direction selectivity. For those neurons, the 5-6 trials at the preferred direction was considered for calculations of ΔF_n_/F0 as the cell’s responses to a particular stimulus. F0 was still averaged over 10-12 trials at two opposite directions.

Several steps were then taken to decide whether each neuron was tuned to orientation and/or spatial frequency. (1) At each spatial frequency, the optimal stimulus size for producing maximal responses at a specific orientation among all orientations was selected. The response to each orientation was decided at this stimulus size. (2) The optimal spatial frequency and orientation producing the maximal response among all conditions were selected. Then responses to 12 orientations were decided at the optimal spatial frequency, and responses to 6 spatial frequencies were decided at the optimal orientation.

(3) To select orientation tuned neurons, a Kruskal-Wallis test was first performed to test whether a neuron’s responses at 12 orientations were significantly different from each other (p < 0.05). For those showing significant difference, the orientation tuning function of each neuron was fitting with a Gaussian model:

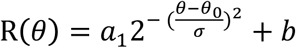

where R(θ) was the response at orientation theta, free parameters a_1_, θ_0_, σ, and b were the amplitude, peak orientation, standard deviation of the Gaussian function, and minimal response of the neuron, respectively. Only neurons with goodness of fit R^2^ > 0.5 were finally selected as orientation tuned neurons.

(4) Similarly, to select spatial frequency tuned neurons, a Kruskal-Wallis test decided whether a neuron responded differently to different spatial frequencies (p < 0.05), and data fitting with a Different-of-Gaussian model further filtered-in neurons with R^2^ > 0.5 as spatial frequency tuned neurons.

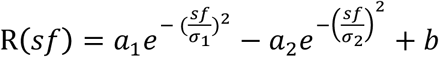

where R(sf) was a neuron’s response at spatial frequency sf, free parameters a_1_, σ_1_, a_2_, and σ_2_ were amplitudes and standard deviations of two Gaussians, respectively, and b was the minimal response among 6 spatial frequencies.

### Optical imaging-like orientation maps

Differential images of 12 orientations were each spatially filtered with a 200-μm low-pass Gaussian filter. Preferred orientation of each pixel on orientation map was then deduced from the vector sum of corresponding response intensities with 12 orientations as following,

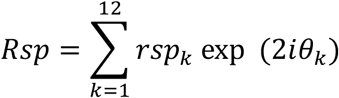

where *θ*_*k*_ was k-th orientation in radians, and *rsp*_*k*_ was the response strength at k-th orientation. Each pixel of the orientation map was then assigned to one of 12 bins according to its orientation preference, denoted by different colors (Figure 3b).

### Autocorrelation and cross-correlation

#### Autocorrelation

Within each orientation or spatial frequency map, every pair of neurons’ correlation coefficient of responses to 12 orientations or 6 spatial frequencies from model fitting were calculated. The correlations of all pairs of neurons separated by a specific range of cortical distance (from 0-800 μm in 50 μm steps) were then averaged as the autocorrelation of this range of cortical distance.

#### Cross-correlation

Same as autocorrelation except neurons were paired from different maps at 150 and 300 μm depths.

## Acknowledgments

This study was supported by Natural Science Foundation of China grants 31230030 and 31730179, as well as funds from Peking-Tsinghua Center for Life Sciences, Peking University. We thank colleagues Geoff Ghose, Stanley Klein, Ming Li, Haidong Lu, Ian Nauhaus, Adam Reeves, and Rufin Vogels for their constructive comments and suggestions during the preparation of this manuscript.

